# Mechanical Analysis of Cell Migration Using Hybrid Structural Optimization

**DOI:** 10.1101/2024.05.03.592447

**Authors:** Eric Havenhill, Soham Ghosh

## Abstract

Cell migration requires the dynamic formation and dissolution of mechanical structures inside the cytoplasm. Stress fibers are made of F-actin during cell migration driven by the strategic localization of focal adhesion complexes at the cell-substrate interface. The nucleus is also strategically positioned in the cell during the migration and the stress fibers wrap around the nucleus possibly to carry the nucleus with the cell. Cell migration is energetically demanding and should require strategic utilization of resources such as the F-actin stress fiber formation at specific locations so that they generate enough force by actomyosin contraction at the cell-matrix adhesion sites for a directed movement. In this work we propose a structural optimization based biophysical model to predict the strategic localization and sizes of F-actin fibers that supports the nucleus and the cytoplasm during migration. With the use of a nonlinear controller via a Newton-Euler-based model of the generated design, we further quantified the force in the stress fibers during migration, with results close to those obtained through experimental methods such as traction force microscopy. The predicted force decreases for a cell that migrates slowly due to a pharmacological perturbation. Such quantification of forces only require the information of the trajectory of the cell that can be readily obtained from time lapse microscopy. With novel microscopy techniques emerging, such biophysical model framework can be combined with traction force microscopy data to achieve unprecedented mechanical information inside and outside cells during migration, which is otherwise not possible by experiments only.

**SIGNIFICANCE:** Cell migration plays a critical role in biological functions. It requires the strategic formation of F-actin stress fibers at specific locations, to generate forces by actomyosin contraction for cells to migrate in a directed manner. The present study predicts the localization and force generated by stress fibers based on the trajectory of the cell, which can be obtained via time lapse microscopy. The technique can complement other techniques such as traction force microscopy to provide mechanical information inside and outside cells during cell migration.

## INTRODUCTION

Cell migration is a fundamental biological phenomenon. Prokaryotes migrate as a single organism to find resources and in this process the cell-cell communication might be critical if the cell is the part of a colony. In eukaryotes, cells migrate to achieve a systemic goal such as tissue formation and regeneration. Immune cells migrate to rescue tissues from infection and cancer cells migrate to metastasize from a primary location to a secondary location (1, 2). In some situations cells migrate alone and based upon the need, cells migrate together showing some specific attributes of collective cell migration where mechanical and chemical communication between cells occur (3, 4). For example, a few leader cells are shown to drive the follower cells in the collective cell migration and the follower cells take turn to be the leaders when the previous leaders run out of the energy (5, 6). Mechanically, cells can communicate with each other via the compliant matrix thus having possible effects in the collective cell behavior (7), such as the speed of individual cell migration. Traction force generated by cells are shown to be different when cells are away from each other compared to when they are closer (8), thus possibly contributing to cell migration by cell-cell mechanical communication.

Structures inside the cell such as F-acin protein in the cytoskeleton and the focal adhesion complex at the cell-matrix interface are critical for the energetically demanding cell migration process. On an individual level, cell movement can be divided into a few stages: protrusion of the leading edge of the cell, adhesion of the leading edge with the matrix (in 3D) or substrate (in 2D), release of the cell from the matrix or the subsrate at the cell body and rear, and cytoskeletal contraction to pull the cell forward (9). The force from the actin polymerization drives the cell membrane forward - the protrusion stage. Subsequently the polymerized F-actin connects to the substrate at focal adhesion points. The contraction, where the bulk of the trailing cytoplasm is drawn forward, is executed by the protein myosin. Thus, the actomyosin contraction and the resultant cytoskeletal forces are mechanically important factors in cell migration. Experimental image of a migrating cell shows such F-actin cytoskeletal structures, called stress fibers Fig. 1. Cytoskeletal forces in the stress fibers are related to the traction force created by the cell at the cell-matrix interface. The quantification of the traction forces during cell migration serve as valuable information for cell mechanosensing and understanding the driving mechanical components to cell migration (10). Because cell migration is energetically demanding requiring the cells continuously forming and dissolving structures, it is anticipated that structures are optimally created inside the cell for most efficient utilization of the resources expending least amount of energy.

**Figure 1.**
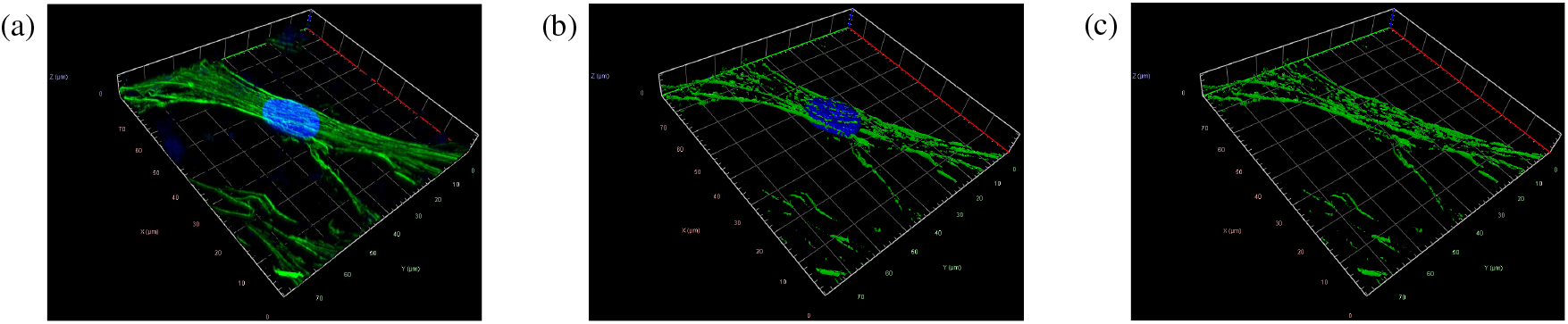
Confocal microscopy images of a migrating murine fibroblast (NIH 3T3 cell). (a) 3D reconstruction of the raw image stack showing the actin (green) and nucleus (blue). (b) Segregated image shows the actin stress fiber with the nucleus and (c) showing the actin stress fibers only.

The nucleus also plays a critical role during the cell migration as recent studies indicate (11). During the migration, the nucleus must be moved with the cell which possibly dictates the location of the centrosome during cell migration (12). The stress fibers that are produced as a part of the dynamic cytoskeleton serve as the structure in ‘carrying’ the nucleus during migration. Previous studies have investigated the stress fiber orientation (13, 14), and different predictive models have been created to study the formation of the stress fibers as the cell moves, as well as the forces that these stress fibers exert on the substrate (15–17). However a critical knowledge gap exists regarding the understanding of cell migration - (1) where the stress fibers are strategically formed that meets the cell’s demand to move forward with the nucleus and (2) what forces those stress fibers create to move the cell forward along with the nucleus? Answering these questions utilizing experiments will require high resolution live microscopy combined with stress calculation inside and outside cells which is currently limited. Biophysical model can be utilized to fill this knowledge gap, which is the primary focus of the present study.

## MATERIALS AND METHODS

### Modeling of a 3 × 3 cell array under resting state using FEM

To quantify the stress distribution in cells a 3 by 3 cell array in two dimensionm was created in FEBio. Tetrahedral elements were used to mesh the combined model of the cytoplasm and the nucleus. The cytoplasm of the cells are made with a neo-Hookean material, with *ρ* = 1*g*/*μm*^3^, *E* = 0.5*k Pa* and *v* = 0.35 for the density, Young’s modulus and Poisson’s ratio, respectively. A prescribed isotropic active contraction was included in the material property, with a contractile stress value of *T*_0_ = 0.01*kPa*. The nuclei of the system are made with a neo-Hookean material, with *ρ* = 1*g*/*μm*^3^, *E* = 1.2*k Pa* and *v* = 0.35 for the density, Young’s modulus and Poisson’s ratio, respectively. The substrate of the system is made with a neo-Hookean material, with *ρ* = 1*g*/*μm*^3^, *E* = 5*k Pa* and *v* = 0.35 for the density, Young’s modulus and Poisson’s ratio, respectively.

### Methodology to quantify the distribution of cytoskeletal material in cell

Modern optimization theory has revolutionized various engineering disciplines by enhancing capabilities in tasks such as generative design of structural systems for improved efficiency, such as minimizing mass while maintaining structural integrity, and facilitating dynamic path planning for enhanced system performance. Utilizing calculus of variations, both static and dynamic optimization problems are commonly formulated. These problems can be addressed through different methods, namely indirect, direct, and hybrid approaches. Indirect methods aim to establish and solve the necessary and sufficient conditions analytically, which is suitable for simpler systems but becomes complex rapidly. Hybrid methods, on the other hand, numerically solve these conditions, often employing shooting methods. Direct methods eschew formulating the necessary and sufficient conditions, instead opting to discretize the system and employ minimization algorithms for solution-seeking. In this paper’s optimization implementation, direct methods are employed.

To begin the optimization, the problem will first need to be posed within a specific framework. Eq. 1 shows an example of how fully dynamic optimization problems are typically posed (18). The cost J to be minimized is the Bolza cost that includes the Mayer term,Φ(·), and the Lagrange term, L(·), which is subjected to constraints on the system, where ***ε***(·), ***β***(·), and ***Π***(·) are the vectors that represent the equations of equilibrium, also referred to as the dynamics, boundary constraints, and path constraints, respectively. Contrast ***ε*** with ***ε***, where the latter represents the strain tensor.

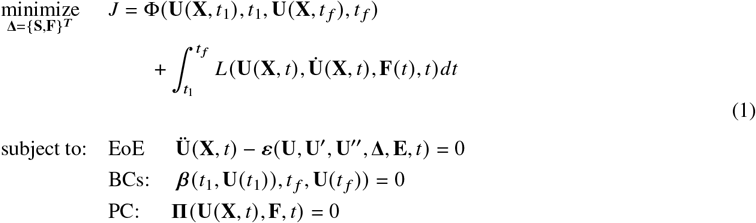

In Eq. 1 for the case of the 1D stress fibers the vector **U**(**X**, *t*) ∈ ℝ^2×1^ is the vector representing the displacement field of the system, as shown in Eq. 2, where **X** is the vector containing the global coordinates of the system, i.e. **X** = {*X, Y, Z*}^*T*^. Contrast the **U** with **u**^(*p*)^ ∈ ℝ^2×1^ which represents the local displacement of the p_*th*_ stress fiber of an element in the finite element method (FEM) - based construction. The vector **x**_(*p*)_ ∈ ℝ^2×1^ represents the spatial coordinates expressed in the local coordinate system of the p_*th*_ stress fiber.

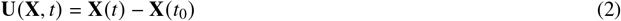

The vector **F** ∈ ℝ^2×1^ represents is the open-loop control, which in this context is a distributed load that causes the contractile or tensile stress fiber forces from the chemical reaction of the nucleotide adenosine triphosphate (ATP). The vector **f** _(*p*)_ ∈ ℝ^2×1^ is reserved for the distributed load across the *p*^*th*^ stress fiber. **S** ∈ ℝ^2*p*×1^ is the vector containing a certain set of states of the system; for this paper, these states are the cross-sectional areas and lengths of the stress fibers, grouped as defined in Eq. 3.

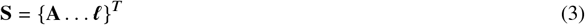

**A** ∈ ℝ^*p*×1^ is the vector that contains all *p* cross-sectional areas of the stress fibers. The vector ***ℓ*** ∈ ℝ^*p*×1^ contains the lengths of the stress fibers. The vector **E** ∈ ℝ^*p*×1^ represents the modulus of elasticity of all *p* stress fibers. In this paper, the cost in Eq. 1 will be proportional to the collective mass of the stress fibers, specifically the volume of the stress fibers, due to the finite resources from the biological limitations of the cell. The volume is formulated as the product of the length and cross-sectional areas of the stress fibers. The vector **Δ** represents the parameters of the system that can be adjusted in order to minimize the value of *j*, in the literature called *decision variables*, which are formulated depending on the context of the problem.

### Static Structural Optimization

The equations of equilibrium in Eq. 1 take the form of a partial differential equation (PDE), specifically in a structure similar to the wave equation, however for some systems, such as the ones presented here in Eq. 4 and later in the Newton-Euler model, the equation of equilibrium is an ordinary differential equation (ODE), rather than a PDE. The full equation of motion for an axial bar is given in Eq. 4, where *A, E*, and *ρ* represent the cross-sectional area, modulus of elasticity, and mass-density of the stress fibers; *f* represents a distributed load on the stress fiber. This equation establishes the relationship that the forces in the stress fibers have on the remaining decision variables.

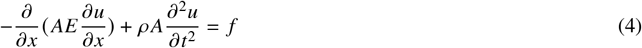

For statically-modeled systems, the time-dependent terms, such as **Ü** or **ü**, will not appear in the static formulations, however the spatially-dependent terms will still appear. Static structural optimization problems are commonly posed in some variation of the way shown in Eq. 1, where the Lagrange term is left null and the Mayer term is commonly chosen as a function that represents the compliance, so as to maximize the stiffness, commonly to minimize the displacement of the structure.

### One-Dimensional Elements

The cellular system of consideration with stress fibers consists of one-dimensional elements, no cytoplasm contribution, and modeling the nucleus as a point mass is illustrated in Fig. 3, where both the protruding and non-protruding stances are considered.

The optimization problem is posed as shown in Eq. 5, where the objective is to minimize the collective mass of the stress fibers. Note that the lengths of the stress fibers are not included in **S** since this is a static optimization problem and hence the lengths are fixed. It is assumed that the same value for the maximum allowable stress is the same in each stress fiber. The governing differential equation for the displacement of an axial bar is given in the constraints. Buckling constraints are also considered in this model, where the inequality constrains the axial load in the bar to be less than the critical buckling load. Note that the lowest modeshape of buckling is considered due to the fact that the fibroblasts move very slowly and as such the higher frequency buckling modes will not be expected at any point. The boundary conditions (BCs) are Dirichlet and defined such that the displacements at the nodes located on the ECM are zero.

**Table 1:** The values of the parameters of the simplified model in Fig. 3.

|  | SF<sub>p</sub> | θ<sub>p</sub> | L<sub>p</sub> (μm) | E<sub>p</sub> (MPa) | A<sub>p</sub><sup>*</sup> (μm<sup>2</sup>) | E<sub>p</sub> (MPa) | A<sub>p</sub><sup>*</sup> (μm<sup>2</sup>) | E<sub>p</sub> (MPa) | A<sub>p</sub><sup>*</sup> (μm<sup>2</sup>) |
| --- | --- | --- | --- | --- | --- | --- | --- | --- | --- |
| Symmetric |  |  |  |  |  |  |  |  |  |
|  | 1 | 15° | ≈ 100.5 |  | 4.5 |  | 4.5 |  | 4.5 |
|  | 2 | 40° | ≈ 40.4 |  | 0.75 |  | 0.75 |  | 0.75 |
|  | 3 | 65° | ≈ 28.7 |  | 0.75 |  | 0.75 |  | 0.75 |
|  | 4 | 90° | 26 | 0.5 | 3.160 | 1.45 | 3.14 | 2.4 | 3.135 |
|  | 5 | 115° | ≈ 28.7 |  | 0.75 |  | 0.75 |  | 0.75 |
|  | 6 | 140° | ≈ 40.4 |  | 0.75 |  | 0.75 |  | 0.75 |
|  | 7 | 165° | ≈ 100.5 |  | 0.45 |  | 4.5 |  | 4.5 |
| Protruding |  |  |  |  |  |  |  |  |  |
|  | 1 | 90° | 26 |  | 3.143 |  | 3.142 |  | 3.140 |
|  | 2 | 102.5° | ≈ 26.6 |  | 0.75 |  | 0.75 |  | 0.75 |
|  | 3 | 115° | ≈ 28.7 |  | 0.75 |  | 0.75 |  | 0.75 |
|  | 4 | 127° | ≈ 32.8 | 0.5 | 0.75 | 1.45 | 0.75 | 2.4 | 0.75 |
|  | 5 | 140° | ≈ 40.4 |  | 0.75 |  | 0.75 |  | 0.75 |
|  | 6 | 152° | ≈ 56.3 |  | 0.45 |  | 4.5 |  | 4.491 |
|  | 7 | 165° | ≈ 100.5 |  | 0.45 |  | 4.5 |  | 4.4499 |

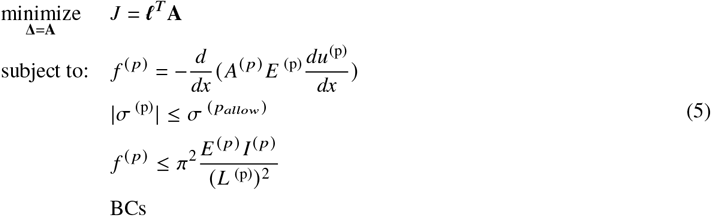

### Three-Dimensional Elements

For a higher-dimensional model, particularly in three dimensions, the structural optimization problem can be posed in Eq. 6, where J is the cost function to be minimized and m is the mass of the system. The governing equations of equilibrium, stress-strain relations, and strain-displacement relations that act as a constraint on the system are expressed in indicial form, where i, j, and k correspond to the x, y, and z components. For a planar model, i = 1,2 and j = 1,2. For a three-dimensional model, i = 1,2,3 and j = 1,2,3.

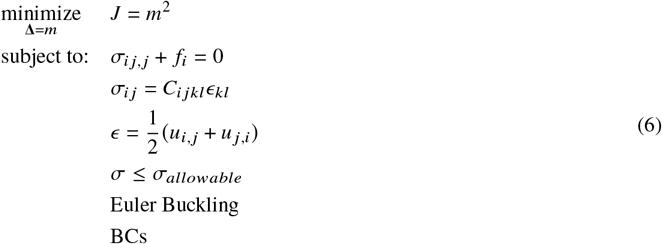

### FEM-Based Transcription Method

The formulations presented in Eq. 1, and hence Eqs. 5 and 6, are continuous-time formulations, which means that they will have to be converted into an approximate discrete formulation for the direct collocation through transcription into a nonlinear program, similarly to the process in (19). The transcription will result in a system of equations that take the structure of Eq. 7, where ℳ, 𝒦, 𝒢, and ℱ represent the mass and stiffness matrices, respectively, and the gravitational force and other force vectors, respectively.

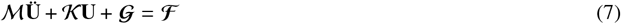

Using this method, several solutions to the structural optimization are obtained and included in this paper. Here, the structural optimization was solved in MATLAB with the use of ‘fmincon’.

### 1D FEM-Based Transcription Method

Only the intermediate region of the cell that is bounded by the nucleus and the cell membrane, and not the nucleus was optimized. The connection of the stress fibers (SF) to the extra-cellular matrix (ECM) and the nucleus is modeled with a pin-joint connection. The connection of the stress fibers to the nucleus is modeled at the center of the nucleus for simplicity. The goal was to minimize the total mass of the stress fibers. This optimization formulation is chosen because the cell has finite resources with which to construct the actin stress fibers, hence we should expect the cell to seek the usage of the minimum mass possible for the construction of each stress fiber.

The equations in Eq. 5 were re-expressed, through transcription, before the optimal solution was obtained with MATLAB’s fmincon. This process involves the converting of the ODE into an approximately equivalent set of algebraic equations, which can be done with several methods; here, the finite element method is used. For a small system like this, the stiffness matrix can be easily obtained and assembled; the building block for each stress fiber’s contribution is given in Eq. 20. It was assumed that the modulus of elasticity will be the same for all of the stress fibers. Note that this matrix is formulated from the use of linear shape functions. Complete details of the finite element-based transcription are reported later.

The reduced matrix, which is obtained upon applying the boundary conditions of zero displacements at all of the nodes located at the substrate side, was used in Eq. 8. The buckling criteria was re-expressed in terms of the displacements and cross-sectional areas. The stress constraints was re-expressed in terms of the displacements. Note that |σ_*p*_ | ≤ σ_*allow*_ is mathematically the same as *−*σ_*allow*_ ≤ σ_*p*_ ≤ σ_*allow*_; the latter formation was used so that the stress constraint could be inserted into fmincon. Values for the material properties, particularly the breaking force, cross-sectional radius, and the modulus of elasticity of the stress fibers were assumed as 377 nN, 100 nm, 1.45 MPa, respectively (20). Therefore, the ultimate tensile strength of the stress fibers was assumed to be 12 MPa. The stress fiber length could vary in length, from 10 to 100 *μm*; typical stress fiber lengths are around 100 *μm* (21). Also, the stress fiber length was ≥ 20*μm* (14). The cross-sectional area of the stress fibers can range from 0.001 *μm*^2^ to 0.04 *μm*^2^, whereas the focal adhesion cross-sectional areas can vary from approximately 0.75 *μm*^2^ to approximately 4.5 *μm*^2^(21). Values of the traction forces during cell motion range from roughly 0 nN to roughly 120 nN (10); 120 nN was used in the simulation. The optimization problem posed in Eq. 5 was converted into a roughly equivalent optimization problem, shown in Eq. 8.

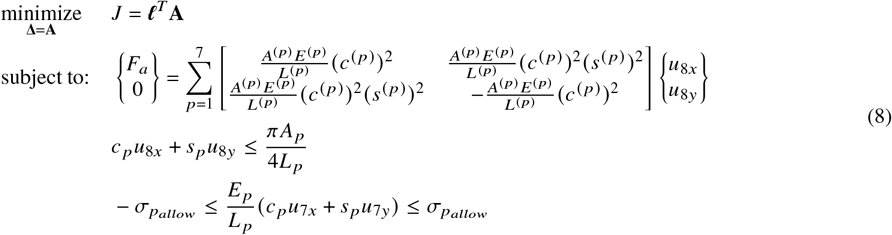

### FEM Formulation - 1D Element

A numerical solution was obtained with the use of the finite element method. The weak form was obtained as shown in Eq. 9, where *ζ* (*x*) represents the weighting function. Note that the superscript denoting the *p*^*th*^ element is not shown; the finite element formulation is the same for each stress fiber, so the superscript here is omitted.

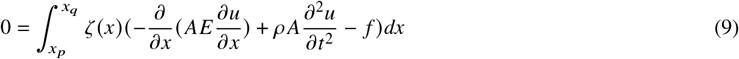

Regardless of the order of the time derivative, there is no integration by parts in time. The final expression for the semi-discrete weak form is shown in Eq. 10.

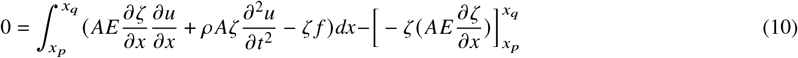

The displacements and weighted residuals are defined in terms of the shape functions as shown in Eqs. 11 and 12.

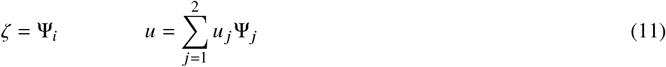

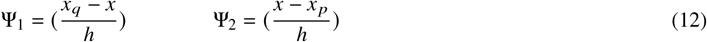

Inserting these equations into the former expression for the semi-discrete weak form will result in the system of ODEs in Eq.13.

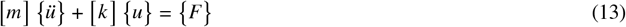

The values of the entries of the mass matrix, stiffness matrix, and force vector are given in Eqs. 14-16.

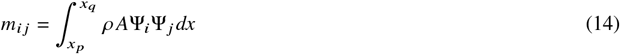

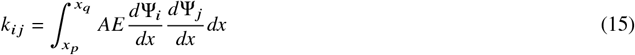

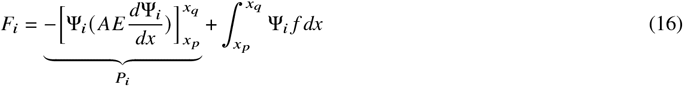

The system was converted using the parent-coordinate formulation in order to formulate the mass matrix. This is shown for the mass matrix in Eqs. 17 and 18.

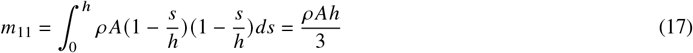

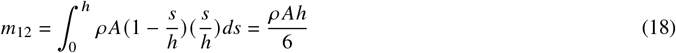

Assembling this results in the mass matrix for a single element as shown in Eq. 19.

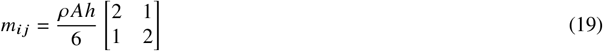

### FEM Formulation - Rotated Element

For the purposes of the combined stress fiber/nucleus system, the finite element formulations were expressed in the global X, Y, Z system, as opposed to their local coordinate systems, in order to formulate the global mass and stiffness matrices as well as the global gravitational and force vectors. Performing the coordinate system transformation resulted in the system of equations for the static case shown in Eq. 20, where *m* and *n* represent the left and right nodes on the one-dimensional finite element, respectively. The stiffness matrix of the rotated bar element is *k* _*p*_ and *c* _*p*_ = *cos*(*θ* _*p*_), *s* _*p*_ = *sin*(*θ* _*p*_).

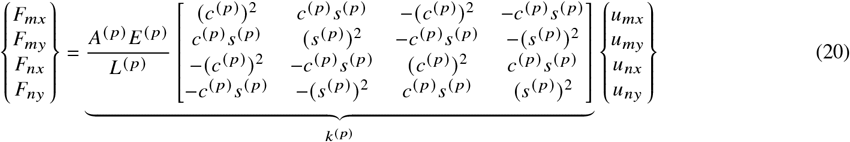

### FEM Formulation - Combined System

The finite element formulations for each element are used to assemble the reduced global system of equations in a succinct manner that arise from the application of the boundary conditions. The matrix ℳ in Eq. 21 is known as the reduced mass matrix of the discretized system, which is full rank and invertible and obtained after applying the boundary conditions. This is contrasted with the global mass matrix, which is not full rank and hence not invertible.

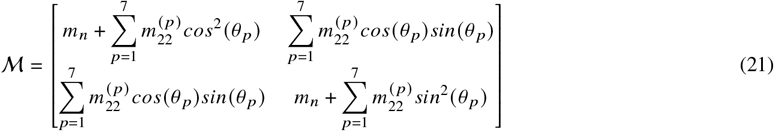

The matrix **𝒦** in Eq. 22 is known as the reduced stiffness matrix of the discretized system, which like the mass matrix, is full rank, invertible, and obtained after applying the boundary conditions. Like the mass matrix, this can be seen in contrast with the global stiffness matrix, which is not full rank and not invertible.

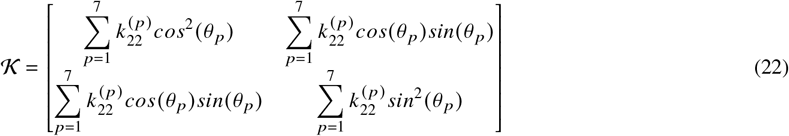

The vector **𝒢** in Eq. 23 represents the contribution due to gravitational forces that arise when considering gravity in the vertical direction with the nucleus’ being modeled as a point mass at the end of the stress fibers, where *m*_*n*_ represents the mass of the nucleus.

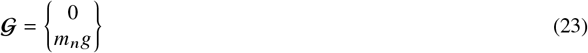

Lastly, the **ℱ** in Eq. 24 represents the forces produced by the stress fibers that support the nucleus and carry the nucleus along the desired trajectory.

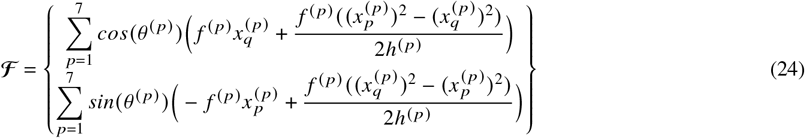

### 3D FEM-Based Transcription Method

The optimization problem posed in Eq. 6 can be converted into an approximately equivalent form, by expressing the constraint equations as a system of algebraic equations; one possible way to do this is through the finite element method. By applying the boundary conditions, the reduced matrix system, with corresponding stiffness matrix K, can be obtained. Subsequently, the optimization problem can be posed in Eq. 25, where **U, V**, and 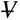 represent the global horizontal displacement, vertical displacements, and volume of the system, respectively.

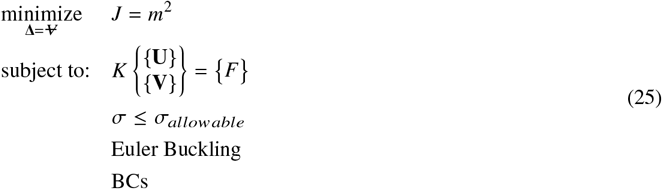

## RESULTS

### Finite element method predicts significantly different stress distribution in the cell based on cell-substrate contact model

Mechanical stress and strain in the cell can be modeled utilizing the Finite Element Method based techniques. Previous works modeled such mechanical quantities in a resting cell utilizing a material model in the cell and the nucleus, along with a suitable boundary condition such as the traction forces (22). One approach is to define a traction force field as a continuous field variable at the cell surface. A second approach is to define the traction force at individual points where the cell matrix adhesion is formed. Traction force is the result of cell contractile force which is not common in modeling cell mechanics but can capture more fundamental aspects of the physical forces in cells compared to the traction force. In the present work we applied a contractile stress inside cells to model the resting mechanics inside the cells using FEBio, a nonlinear finite element analysis software created specifically for biomechanics and biophysics applications (23). The goal is to investigate the stress distribution inside cells.

In our model we incorporated the provisions to investigate the stress and strains inside the cell, the nucleus and the substrate in the three-dimensional setting. Nine cells were arranged in a 3 × 3 array with the distance between cells being a possible parameter that can be changed. For visualization purpose only a planar slice is shown 2. In the first case (a), continuous surface contact was assumed and in the second case (b), specific contact areas were modeled utilizing ‘feet’ under the cells to mimic the focal adhesion. The results reveal that when the “feet” are included, the stress is concentrated largely at the end regions of the cell, with smaller amount of stress in the middle, as opposed to the continuous distribution observed when the cell makes a continuous flush contact with the substrate.

Multiple simulations were performed, where the contractile stress *T*_0_ inside the cell was perturbed in order to investigate the effect of different contractile stress values on the stress magnitude field shown in the color plot in Fig. 2. Such parametric study changes the magnitude of the stress field inside the cell, but it does not change the shape of the stress field, therefore results from multiple simulations are not included since the purpose of this particular model is to look at the shape of the stress distribution that arises from the cell’s contraction during migration, not the magnitude. Overall this results indicate that only specific areas of the cell experience an elevated amount of stress and they are close to the outer periphery of the cell whereas inner locations undergo lower amount of stress. The F-actin tread milling is a mechanical stress dependent process therefore the stress fibers are most likely formed at those adhesion points near the cell periphery and connecting the cell across with fiber formations as shown in the images from the experimental data in Fig. 1.

**Figure 2.**
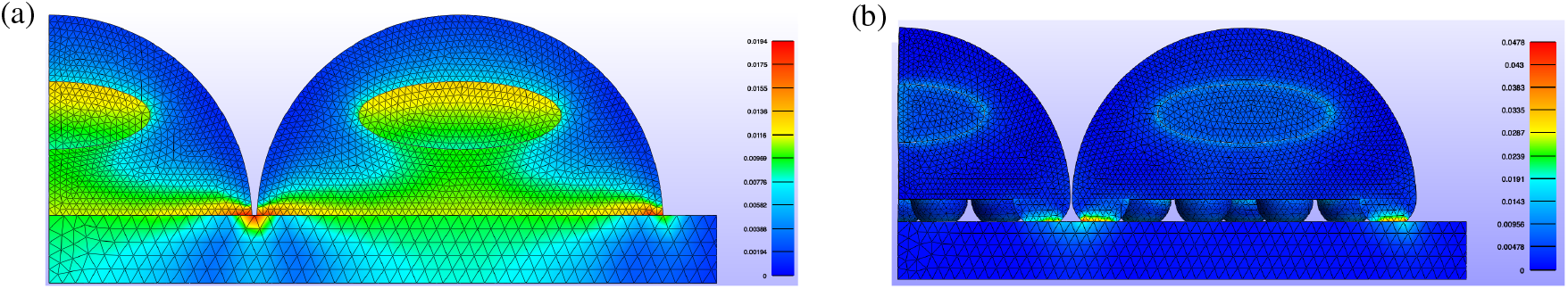
Distribution of ‘stress magnitude’ inside the cell, nucleus and substrate under resting state modeled with cell contractility inside the cytoplasm (a) with the cell modeled under perfect contact with the substrate and (b) with focal adhesion points that contact the substrate at specific locations marked by the ‘feet’.

### Stress fiber thickness is optimally defined by cells at strategic locations during migration as revealed by a planar cell model

A handful of stress fibers are formed during cell migration that build the mechanical structure to carry the cell forward as shown in Fig. 1. The nucleus is connected to the cytoplasm via the F-actin fibers and the nucleus also moves with the cell in a coordinated manner (24). Therefore, it is expected that the F-actin fibers carry the nucleus. This leads us to the question - how only a few stress fibers are mechanically able to carry the nucleus. This is logical though because the F-actin fiber formation is energetically demanding and cells will try to optimize the stress fiber formation at strategic locations and numbers with a goal to carry the cell and the nucleus forward.

Therefore we attempted to predict the thick stress fiber formations at specific focal adhesion site locations by solving an optimization problem that seeks to minimize the collective mass of the stress fibers. The model of the cell is shown in Fig. 3. The top panel shows the configuration of the cell at resting (a) and protruding states (b). The thin lines represent the possible locations of the stress fibers. Subsequently the optimization problem was posed and executed to calculate the requirement and thickness of the fibers, sufficient to carry the nucleus subjected to a force *F*_*a*_. in this case, *F*_*a*_ is chosen to be 120 nN, the magnitude of the force was derived from the literature (10). The numerical values of the lengths of the stress fibers and their orientations are given in Table 1.

**Figure 3.**
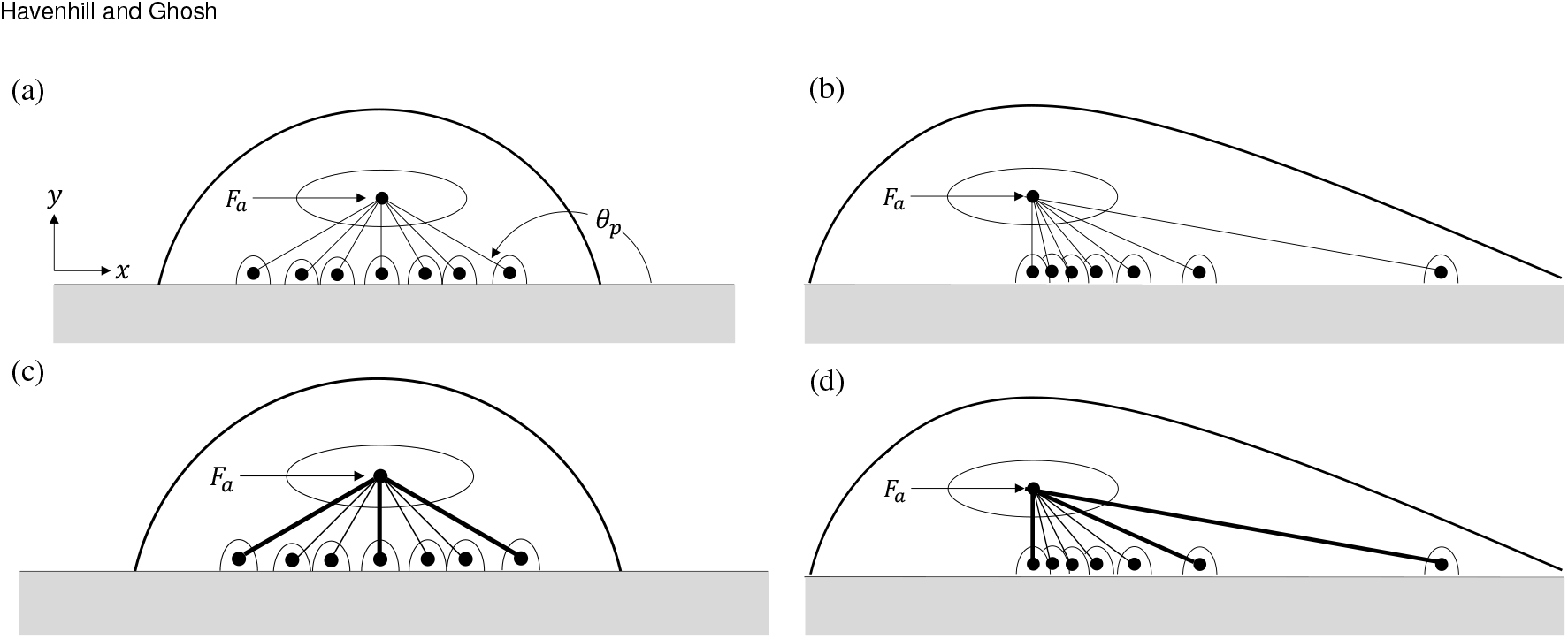
(a) Model of the simplified cell with the possible locations of stress fibers and associated focal adhesion points shown for (a) the symmetric case and (b) the protruding case. The cell with the stress fibers, with thicknesses shown to scale after the static structural optimization, and associated focal adhesion points presented in (c) before protrusion and (d) during protrusion. For complete description of stress fiber areas after optimization refer to Table 1.

MATLAB’s ‘fmincon’ was used in order to optimize the system under such mechanical loading demand which provides the optimal cross-sectional areas of the stress fibers. The values of the areas from the optimization, as well as the parameters of the model, are presented in Table 1. Note that the (·)^∗^ represents the optimal solution. After thresholding, we get that *A*^(2)^ ≈ *A*^(3)^ ≈ *A*^(5)^ ≈ *A*^(6^ ≈ 0, which leaves just the contribution from stress fibers 1,4, and 7. The resulting system is shown in Fig. 3 c and d. The thicknesses of the fibers have been scaled for visualization purposes. It should be noted that *F* _*a*_ can be varied within the neighborhood of this value, however the distribution of the three principal stress fibers remain the same, suggesting that an exact value of this force is not critical. Overall this results agree with the experimental observation that only a few thick stress fibers are formed in the cell to carry the nucleus forward. A three dimensional counterpart of this model is also framed and presented in Fig. 7 in the appendix.

### Prediction of traction forces from the cell trajectory

#### Dynamic model to predict the cell trajectory

The results presented so far were obtained by applying a static structural optimization on the cell’s cytoskeleton. However, a more sophisticated model is desired that accounts for a dynamic system by utilizing the trajectory of the nucleus. The trajectory was used to quantify the position, velocity, and acceleration of the nucleus, so dynamic equations were generated using a technique we previously described (19), that utilizes the dynamic mode decomposition (DMD) method. Equations for such trajectories that represent position, velocity and acceleration are presented below in Eq. 26. The cell (or nucleus) trajectories were derived from our previous experimental work (11), where murine embryonic fibroblast line NIH 3T3 cells were visualized during migration in a scratch wound assay. We utilized the data from an experimental perturbation where a drug Trichostatin-A lowers the cell migration velocity by inhibiting the chromatin condensation (11).

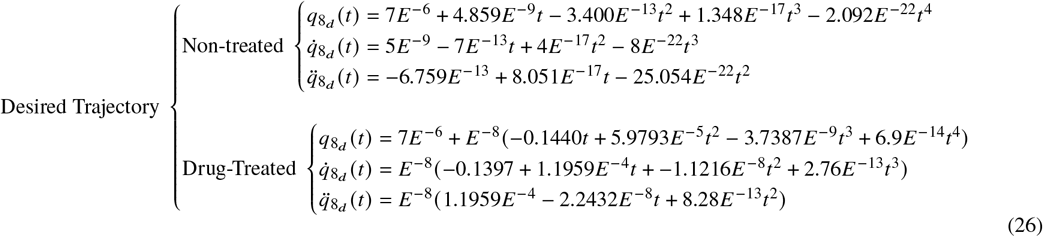

In order to illustrate the resemblance between the experimental data obtained from ImageJ’s TrackMate (25) and the resulting DMD process, the trajectories of both for the control and drug-treated (TSA treated) cells are shown in Fig. 4. Procedures towards developing a desired trajectory that will be used in the control selection can take multiple approaches; here, the DMD-based trajectory is used for it’s higher *C*^*k*^ continuity, where here *k* represents the *k*^*th*^ derivative, compared to the experimental trajectory. A smoother desired trajectory function helps in eliminating any jerky movement of the system as well as it allows the easier formulation of the desired kinematic trajectories.

**Figure 4.**
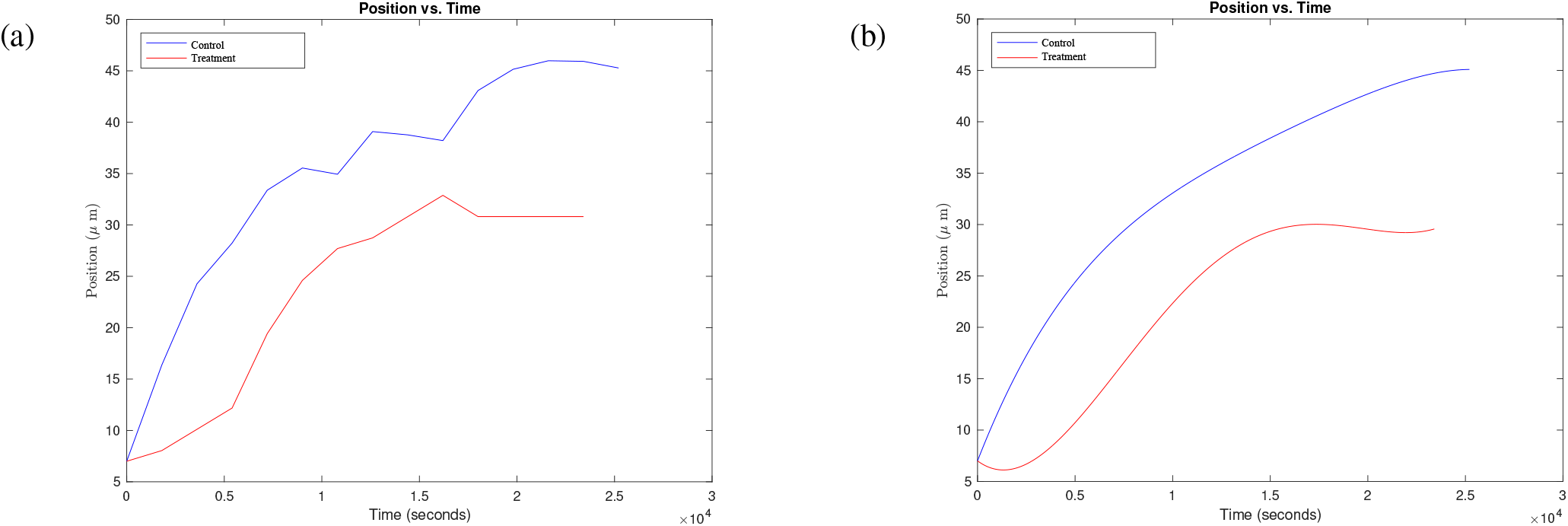
(a) Plot of the experimental data where the starting x-values of both plots have been adjusted to start at 7 *μ*m for trajectory comparison and (b) the plot of the approximated trajectory obtained through dynamic mode decomposition based on the experimental data.

#### Newton-Euler hybrid-based model to represent the stress fibers as robotic elements

One approach towards obtaining a dynamic model is to construct a multi-link Newton-Euler based “robotic-like” system as a model for the cell. The benefit of such an approach includes the ability to apply well-established control theory methods to this system. The “robotic-like” model of the cell is shown in Fig. 5. The coordinate directions are defined by the inertial reference frame **N**, where the orthonormal unit vectors that define the basis of the *right-handed* spatial coordinate system are 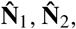, and 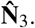.

**Figure 5.**
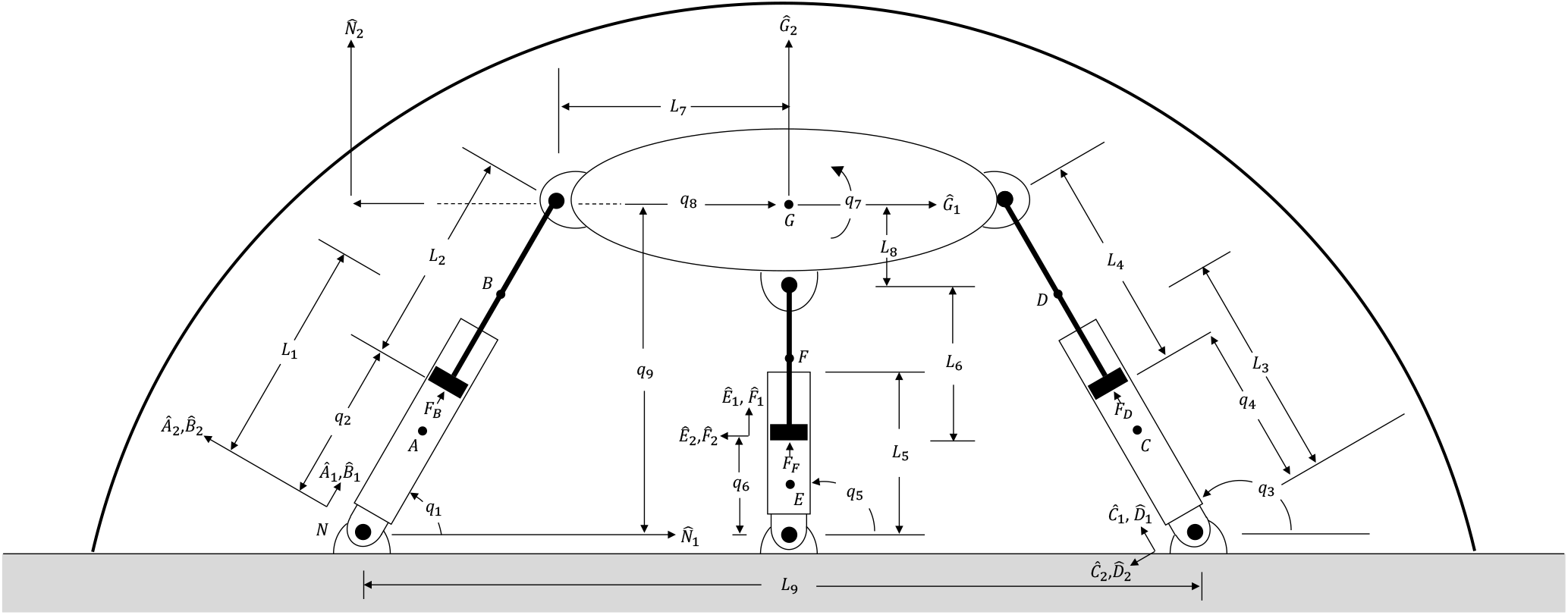
Coordinate system diagram of the “robotic-like” model of the cell with three actuator-based stress fibers. The connections to the actuators at both the substrate and nucleus are modeled as pin joints. The generalized coordinates representing translational or rotational position are represented by *q*_1_…*q*_9_. The body-attached frames for each rigid body are **A** = [*Â*_1_, *Â*_2_, *Â*_3_,], …, **G** = [*Ĝ* _1_, *Ĝ*_2_, *Ĝ*_3_,], where *Â*_1_,…, *Ĝ* represent the orthogonal set of unit vectors defining each frame. Relevant lengths of the rigid bodies are given as *L*_1_,…,*L*_9_. All bodies, including the substrate, are assumed to be completely rigid.

The dimensions of all of the lengths of the rigid bodies in this robotic representation of the cell, as well as all of the initial values of the generalized coordinates, are shown in Table 2, where *L*_*n*_ represents the length of the *n*^*th*^ element and *q*_*n*_ represents the *n*^*th*^ generalized coordinate. The size of the nucleus of a fibroblast ranges between 500 *μm*^3^ to 1000 *μm*^3^ (26); the size of the nucleus in this work is assumed to be 513.13 *μm*^3^. The mass of the nucleus is assumed as 2.29 ng (27); the mass of the cell is assumed to be 22.9 ng.

**Table 2:** The parameters of the three-link “robotic” model of the cell shown in Fig. 5, at its initial states.

| $n$ | $L_n$ | $q_n(t_0)$ |
| --- | --- | --- |
| 1 | $20 \mu\text{m}$ | $90^\circ$ |
| 2 | $20 \mu\text{m}$ | $6 \mu\text{m}$ |
| 3 | $20 \mu\text{m}$ | $\approx 150.58^\circ$ |
| 4 | $20 \mu\text{m}$ | $\approx 32.926 \mu\text{m}$ |
| 5 | $20 \mu\text{m}$ | $\approx 134.90^\circ$ |
| 6 | $20 \mu\text{m}$ | $\approx 12.759 \mu\text{m}$ |
| 7 | $7 \mu\text{m}$ | $0^\circ$ |
| 8 | $2.5 \mu\text{m}$ | $7 \mu\text{m}$ |
| 9 | $60 \mu\text{m}$ | $26 \mu\text{m}$ |

Several classical mechanics methods exist that can produce the governing differential equations of motion, such as with the Newton-Euler equations, Lagrange’s method, D’Alembert’s principle, Kane’s method, etc. Here, Kane’s method was used with the help of Autolev motion analysis software (28) in order to generate the equations of motion. The governing equations for the system are represented in joint space in Eq. 27, and are subject to the kinematic constraints on the system obtained from the development of kinematic chains shown in Eq. 28.

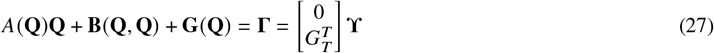

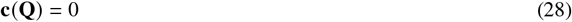

In the equations above, 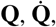, and 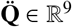 are the vectors containing the generalized coordinates that define the configuration of the system, velocities, and accelerations, respectively, *A*(**Q**) ∈ ℝ^9×9^ is the inertial matrix of the system, **B** ∈ ℝ^9×1^ is a vector containing the Coriolis and centrifugal terms, **G** ∈ ℝ^9×1^ is the matrix containing the terms due to gravity, **Γ** ∈ ℝ^9×1^ is the vector containing the actuator force and torque values, *G*_*T*_ ∈ ℝ^9×3^ is the transmission matrix, and **ϒ** ∈ ℝ^3×1^ is the vector containing only the actuator forces. It is noteworthy that the equations stated above are second-order, nonlinear differential equations that describe the motion of the robotic model of the stress fibers in the cell. The degrees of freedom (DOF), sometimes referred to as mobility, of the system can be determined with the Chebychev–Grübler–Kutzbach criterion in Eq. 29, where *R, j*_1_, and *j*_2_ represent the number of rigid bodies, joints of type 1, and joints of type 2, respectively.

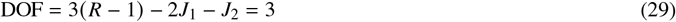

The kinematic constraints on the system, represented by the vector **c**(**Q**) ∈ ℝ^6×1^, are formulated here from vector loops or kinematic chains of the configuration.

#### Nonlinear control of the configuration space equations to quantify the forces in stress fibers

Once the robotic model of the stress fibers were built, we quantified the forces in the stress fibers. To build a classical controller, various methods can be utilized. Here, the equations in Eqs. 27 and 28 were combined through constraint embedding in Autolev and expressed in a single set of equations, as represented in Eq. 30, where 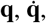, and 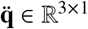 are the vectors containing the generalized coordinates, generalized velocity, and generalized acceleration that remain after the constraint embedding. The matrix **a**(**q**) ∈ ℝ^3×3^ represents the reduced mass matrix, **b**(**q**) ∈ ℝ^3×1^ is the vector containing the Coriolis and centrifugal terms, **g**(**q**) ∈ ℝ^3×1^ is the vector containing the gravitational terms, and 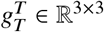 is the reduced transmission matrix.

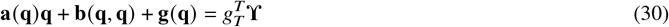

An operational space controller was developed in order to specify the constraint on the movement of the nucleus. Typically, the Jacobian matrix needs to be formulated in order to convert between textcolorredconvert between does not sound right - use some other phrase the joint space and the operational space, if the coordinate of consideration is rotational. However the generalized coordinate representing the nucleus is already expressed in the translational coordinate system, so no Jacobian is needed. The computed torque method was used in order to develop a controller that drives the movement of the system. We selected the following control in order to linearize the dynamics in Eq. 31,

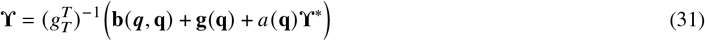

where:

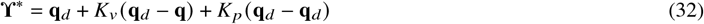

which gives the resulting error dynamics are shown in Eq. 33, where *K*_*p*_ and *K*_*v*_ are the proportional and derivative gains.

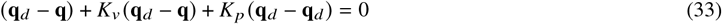

The values of the gains *K*_*p*_ and *K*_*v*_ are selected at the discretion of the user. These gain values will determine if the error dynamics behave in an overdamped, critically damped, or underdamped fashion. The gain selection in Eq. 34 guarantees that the the error dynamics will behave in a critically damped manner; the proportional gain is selected to be *K* _*p*_ = 1. This is a common heuristic model of control that doesn’t necessarily guarantee optimal solution.

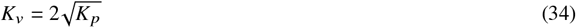

It can be noted that this type of controller can be demonstrated to asymptotically converge the system to the desired trajectory with the use of a Lyupanov-based analysis, even with a variety of gains. To show this, a Lyupanov candidate function for this system is proposed in Eq. 35, where *E* and 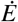 represent the error, defined as *E* ≜ *q*_*d*_ *− q* and time derivative of the error, defined as 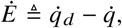, respectively.

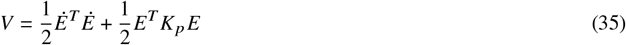

Note that the gain matrix, *K*_*p*_, is a positive-definite matrix. The candidate function V satisfies the criteria for a Lyupanov function, namely V is zero *iff E* and *E* are zero and V is greater than zero *iff E* and *E* are not zero. Differentiating Eq. 35 gives Eq. 36.

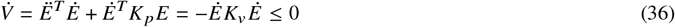

The expression in Eq. 36 is zero only when *E* and 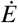 are zero, and negative when *E* and 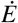 are nonzero, therefore asymptotic convergence of the system to the desired position is guaranteed with the use of any positive-definite gain matrix *K*_*v*_.

The biological insight received from this model is stated below and in Fig. 6. Trajectory of the control cell (that moves faster) and the TSA treated cell (that moves slower) are shown in Figure 6 (a) and (c) respectively. The cells were tracked for almost 7 hours. It is clear that the control cell moved a larger distance along the x axis (straight line) over the same total tracking time. The y component is zero and rotation (*θ*) is also zero because the nucleus was moving in an approximated straight line. The nonlinear controller is able to converge the position (black line) to the desired trajectory (purple line) in both cases, which is consistent with the Lyupanov-based prediction. The force in the individual controllers (stress fibers) change over time but not drastically, which is expected. The distance travelled by the cells in this situation is fairly small - between 30 microns (TSA) to 47 (control) microns which is almost same as the length of cell. Therefore, if we consider a full cycle of cell migration consisting protrusion, contraction and stabilization, the force in individual fibers won’t change much during this timeframe. The comparison of the values of forces in the stress fiber compare remarkably closely with the experimental data derived from traction force microscopy, which reports the traction force in the range of tens of nanonewtons (10).

**Figure 6.**
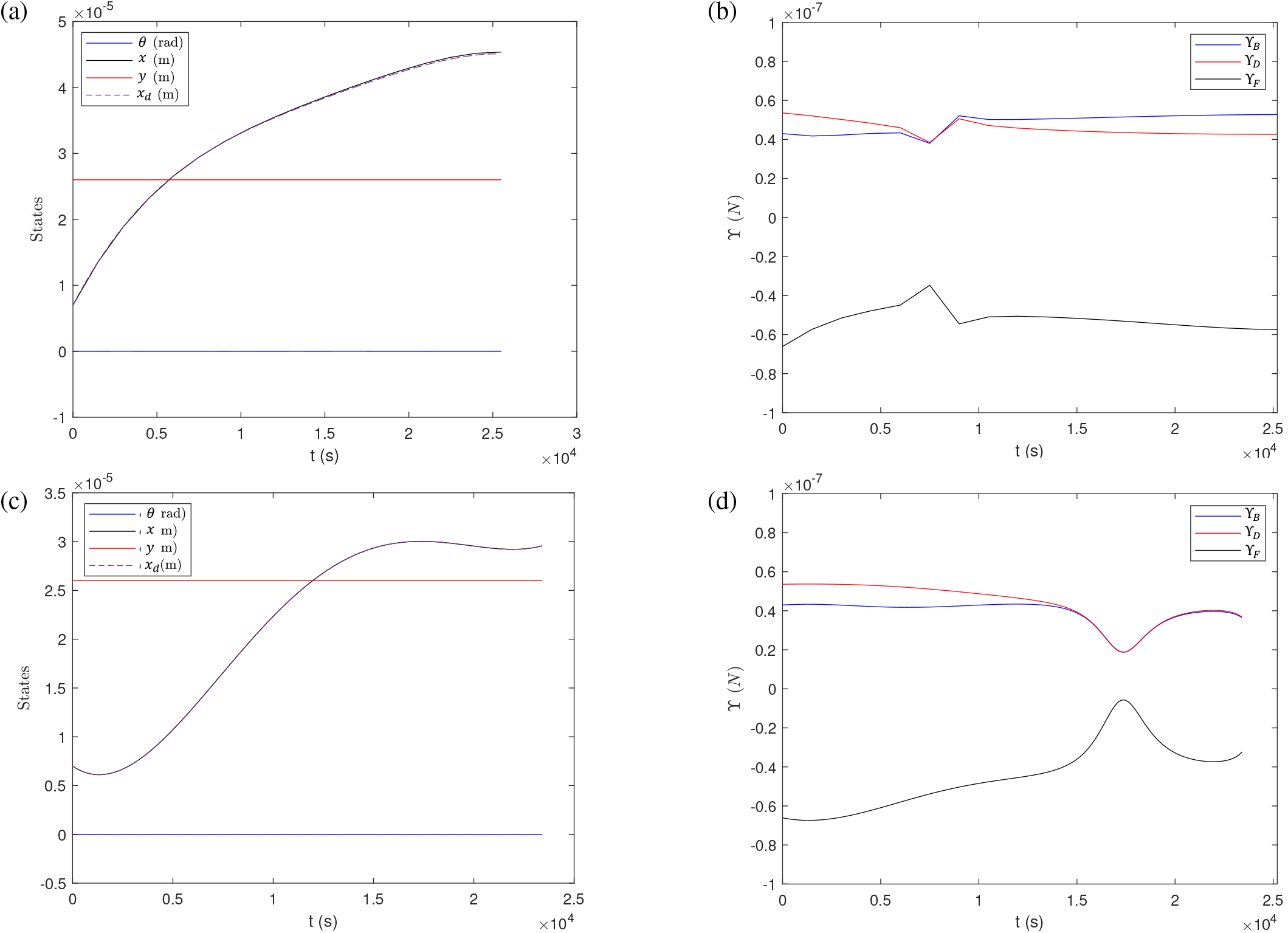
(a) Trajectory of the states of the control cell. (b) Actuator forces from the nonlinear controller of the system for the control cell, where the forces in the left, middle, and right actuators start at 42.97 nN, −66.10 nN, and 53.56 nN; the actuator forces, from left to right, finish at 52.75 nN, −57.41 nN, and 42.62 nN. (c) Trajectories of the states cell with TSA treatment. (d) Actuator forces from the nonlinear controller of the system with TSA treatment, where the forces in the left, middle, and right actuators start at 42.97 nN, −66.10 nN, and 53.56 nN; the actuator forces, from left to right, finish at 36.45 nN, −32.32 nN, and 36.76 nN.

## DISCUSSION

The goal of this research is twofold: to exploit the use of modern optimization techniques in order to predict the orientation and size of F-actin stress fibers networks within the cytoskeleton and to quantify the forces in the stress fibers during cell migration. The migration event was posed as an optimization problem that allows for the cell to achieve a desired motion in a particular direction, subject to a finite actin mass supply constraint. The assembly of the cell’s internal organization at this level during migration is currently unavailable due to the limitations of even the most cutting edge microscopy. Computationally derived quantification of the stress in fibers, that can provide the information of the traction forces is also achieved using the present framework. Only the cell trajectory is the input in this framework which is experimentally obtainable by time lapse microscopy. This can be especially useful when cells are moving in clusters as in those situations quantification of the traction force at the single cell level will require many assumptions. Therefore, to investigate the cell-to-cell communication via mechanosensing in a dense environment we can use the current framework which only relies on the trajectory tracking, which is much easier using time lapse microscopy. However, it should be noted that the method provided in this study is not meant to replace the traction force microscopy, rather to complement that technique to provide the force information both inside and outside the cell.

The optimal location and size of the stress fibers were revealed in this study as shown in Figs. 2 and 3. Although the study was done on a vertical cross-section of the cell (3) to quantify the size of the stress fiber, it still captures the requirement of the cell to create only a few thick stress fibers which matches the experimental observation. A more detailed three-dimensional model with the same technical framework as shown in Fig. 7 also captures the same biological insight. Additionally, the location of stress fibers are concentrated at the periphery of cells which is captured by the high stress concentration in specific locations as shown in Fig. 2.

The quantification of stress fiber forces not only resemble the previously described experimental data, it also shows that the control cells which move faster has a higher value of stress fiber force compared to the TSA treated cell which moves slower. The computational framework only relied on the cell trajectory that was derived from the experiment and the Dynamic Mode Decomposition. The benefit of such an analytical biophysical model as compared to experimental models (traction force microscopy), is that the parameters of the model can be adjusted based on the cell type to obtain the forces in stress fibers.

The framework presented here also exists for dynamic structural optimization problems, however current models are commonly centered around dynamic optimization from a frequency perspective through the augmentation of a frequency constraint (29), such as the one in Eq. 37, where *K, ω*_*r*_, and *M* represent the global stiffness matrix, *r*^*th*^ eigenvalue, and global stiffness matrix, that imposes an eigenvalue problem-type constraint, in place of the equation of equilibrium in Eq. 1 (30). The solution more closely resembles that of a static optimization problem solved in this paper, since the geometry is not designed to change over time. Further improvement of this framework will allow us to predict the redistribution of the cytoskeletal material during the dynamic process of cell migration

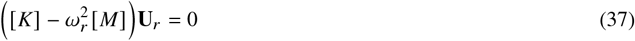

The results presented in this paper set the framework for simulating more sophisticated cell migration behavior. Future work will focus on using the complete equations of equilibrium, as opposed to the reduced static equations, which will allow for a more robust model of cell migration due to the dynamic nature and more elaborate model. This paper also lays the groundwork for allowing more sophisticated three-dimensional model of cytoskeletal material distribution, as presented in Fig. 7. Future work can also add additional constraints to the robotic Newton-Euler model of the cell, specifically by adding frictional or adhesion constraints at the stress fiber attachment points, rather than assuming that these points remain in contact through the entire motion.

**Figure 7.**
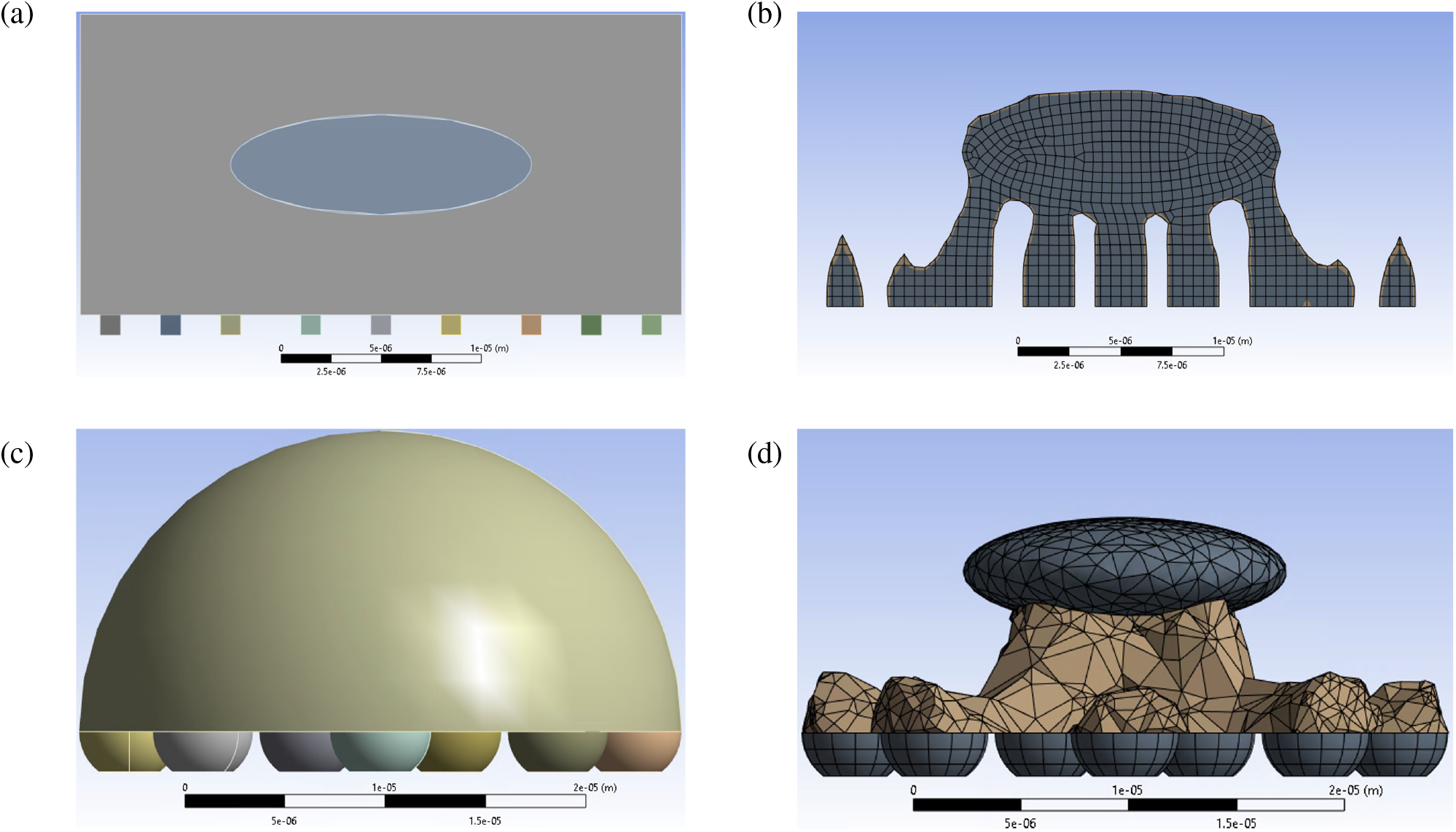
Structural optimizations of the cells in ANSYS are shown. (a) The 2D model of the cell before the optimization. (b) The minimum mass optimization results for the 2D case. (c) The 3D model of the cell before the optimization. (d) The minimum mass optimization results for the 3D case.

## CONCLUSIONS

The outcome of this paper demonstrates a methodology for the creation of a biophysical model that estimates the stress fiber forces and hence it can insinuate the traction forces during cell migration. Additionally it predicts the position and size of the cell’s stress fibers during this process. Therefore, with the power of structural optimization, nonlinear control, and the finite element method, a novel dynamic model of cell migration is achieved, with expected impacts in the field of biomechanics, cell migration and applied optimization sciences.

## AUTHOR CONTRIBUTIONS

E.H. conceived the technical framework, E.H. and S.G. analyzed the results. All authors wrote and reviewed the manuscript.

## SUPPLEMENTARY MATERIAL

The code used in this paper is available via Github at https://github.com/EricHavenhill/SSOpt/.

## APPENDIX

### 3D cell model predicts the stress fiber localization and thickness as a material resource available to cells

A more sophisticated and realistic geometry of the cell can be considered by constructing a model in three dimensions with a goal to predict the stress fiber formation in 3D space. A problem like this can be solved with MATLAB’s fmincon as was done for the simplified cell model. However, due to geometric complexity, the structural optimization is performed in Ansys Topology Optimization. The optimization results from Ansys is shown in Fig. 7. The optimization results compare closely to the optimizations that were shown in Fig. 3. This serves as a verification that the optimization results are converging on the appropriate solution.

